# Unpredictable chronic stress does not alter social behavior in zebrafish

**DOI:** 10.64898/2025.12.15.694431

**Authors:** Náthali A. Neves, Matheus Gallas-Lopes, Amanda Patelli-Alves, Daniela V. Müller, Leonardo M. Bastos, Thailana Stahlhofer-Buss, Ana P. Herrmann, Angelo Piato

## Abstract

Stress-related disorders encompass diverse behavioral alterations, including impaired social functioning. The zebrafish (*Danio rerio*) is a valuable model for studying these phenomena, particularly because of its robust and ethologically conserved social behaviors. Unpredictable chronic stress (UCS) produces behavioral, physiological, and molecular changes in zebrafish that parallel core features of human anxiety and depressive disorders; however, its impact on social outcomes remains unclear. Here, we examined whether a 14-day UCS protocol alters social behavior in adult zebrafish using two complementary assays: the social preference test and the shoal cohesion test. Across two independent experiments, UCS did not elicit detectable changes in individual social approach or group-level cohesion. In contrast, UCS induced clear anxiety-like behavior in the novel tank test, validating the stress manipulation, with stressed fish displaying reduced vertical exploration and decreased time in the upper zone. Shoal cohesion measures showed a time-dependent decrease in both groups, consistent with habituation to the testing environment rather than a stress-specific effect. Together, these results suggest that social behavior in adult zebrafish is relatively resilient to UCS under the conditions tested, whereas anxiety-like responses are markedly affected. Future work should investigate whether factors such as stressor intensity, developmental stage, sex composition, or social hierarchy modulate the sensitivity of social behavior to chronic stress.

## INTRODUCTION

Social dysfunction is a well-recognized symptom of both anxiety and depressive disorders and has major clinical relevance [1]. These impairments often manifest as reduced social engagement, diminished verbal and non-verbal communication, and difficulties in initiating, maintaining, or interpreting social relationships. Individuals with depression may withdraw from social contact, display reduced affective expression, or experience feelings of worthlessness that undermine their social confidence. Similarly, individuals with anxiety disorders, particularly social anxiety disorder, may avoid social situations because of an intense fear of negative evaluation or embarrassment, which can lead to chronic isolation. These disturbances compromise quality of life and social functioning and can contribute to the chronicity and exacerbation of psychiatric symptoms [2–4]. Furthermore, impaired social functioning is frequently resistant to treatment and may continue even after symptomatic remission, highlighting its relevance as both a clinical target and a prognostic factor [5]. Despite their clinical relevance, social deficits remain insufficiently investigated in anxious, depressed, comorbid, and remitted populations [6].

Because stress is a major driver of both anxiety and depressive disorders and a key contributor to the emergence of social dysfunction, preclinical stress models have become essential tools for investigating how chronic stress alters behavior [7]. The unpredictable chronic stress (UCS) paradigm was originally developed in rodent models to mimic the cumulative and unpredictable nature of daily stressors that contribute to stress-related psychopathology in humans [8]. Over several decades, UCS has consistently been shown to induce behavioral, physiological, and endocrine alterations that parallel key features of anxiety and depressive disorders, including anhedonia, heightened anxiety-like responses, cognitive impairments, and dysregulation of the hypothalamic–pituitary–adrenal axis [9]. Building on this framework, UCS procedures were adapted for zebrafish (*Danio rerio*), enabling comparative and high-throughput approaches to the study of chronic stress [10].

Although social behavior in zebrafish is robust and evolutionarily conserved, its sensitivity to chronic stress remains unclear. A systematic review and meta-analysis of UCS studies in zebrafish highlighted that, although outcomes related to anxiety, locomotion, and cortisol generally showed consistent stress-induced alterations, measures of social behavior were highly heterogeneous, with studies reporting reduced, unchanged, or even increased social interaction following UCS exposure [11]. Moreover, relatively few studies have examined social and anxiety-like outcomes within the same individuals, limiting the ability to determine whether UCS selectively affects certain behavioral domains while sparing others. These gaps underscore the need for controlled investigations focusing specifically on social behavior under UCS.

Therefore, the present study evaluated the effects of a 14-day UCS protocol on two complementary measures of social behavior in adult zebrafish: the social preference test (SPT) and the shoal cohesion test (SCT). Because anxiety-like behavior is a well-characterized and reproducible outcome following UCS in zebrafish, in a second experiment we assessed social behavior and performance in the novel tank test (NTT) in the same animals as a behavioral validation of the stress manipulation.

## MATERIALS AND METHODS

This study complied with ethical standards and received approval from the animal ethics committee of the Universidade Federal do Rio Grande do Sul (approval: #42179/2022).

### Animals, housing, and husbandry

We used 216 adult wild-type zebrafish, 3–4 months old, of both sexes, across two experiments: 144 animals in Experiment I and 72 in Experiment II. Because the experimental unit in the SPT is the individual animal, whereas in the SCT it is the shoal (groups of four), we report samples by assay and group composition. Within each experiment, SPT and SCT used non-overlapping sets of animals. In Experiment I, 80 fish (47 males, 33 females) were used in the SPT (n = 40 per experimental group), while 16 shoals (64 fish total; 41 males, 23 females) were used in the SCT (n = 8 shoals per experimental group). In Experiment II, animals were submitted to the SPT (n = 20 per experimental group) or to the SCT (n = 4 shoals per group); all fish were subsequently evaluated in the NTT on the following day. Because sex was determined only after the NTT, sex composition is reported for the NTT: 72 animals, consisting of 33 males and 39 females. The sex of each individual animal, including the sex composition of all shoals, is fully detailed in the raw data files [12]. The study was not designed or powered to test sex differences, so data from females and males were pooled for all analyses.

Animals were obtained from a commercial supplier (Delphis, Porto Alegre, Brazil) and maintained in the zebrafish facility of the Instituto de Ciências Básicas da Saúde (Universidade Federal do Rio Grande do Sul). Rooms followed a 14:10-h light-dark cycle with lights on at 07:00 am. Before behavioral testing, fish were acclimated for at least two weeks in a recirculating rack system (Altamar, Jacareí, SP, Brazil). Fish were kept in unenriched 16-L glass tanks (40 × 20 × 24 cm) with dechlorinated water at a maximum density of 2 fish/L, under continuous aeration and multi-stage filtration. Water quality was maintained within standard zebrafish ranges: 27 ± 2 °C, pH 6.8 ± 0.3, and conductivity 500–800 μS/cm. Animals were fed twice daily with commercial flake food (Poytara®, Araraquara, SP, Brazil) and *Artemia salina*.

During the experimental phase, fish were moved from the recirculating rack to static glass tanks of the same dimensions (16 L; 40 × 20 × 24 cm). Tanks were filled with conditioned water from the same source and held at the same parameters described above. To minimize tank-specific effects, each experimental group was distributed across two tanks (two control tanks and two stress tanks). Tank positions were randomized within the facility so that stress and control tanks experienced equivalent lighting, temperature, and vertical level. Husbandry procedures were matched across all tanks, and opaque barriers prevented visual contact between groups.

At the conclusion of the experiments, fish were euthanized by hypothermic shock (immersion in 2–4 °C water for at least five minutes until opercular movements ceased) followed by decapitation to ensure death. Sex was confirmed by post-mortem gonadal examination.

### Experimental design

The experimental design is illustrated in Figure 1. Experiment I focused exclusively on social outcomes. All animals were evaluated in the SPT or SCT on the morning following the UCS protocol, providing measures of individual social approach and group-level cohesion. Based on the results from Experiment I, we designed Experiment II to replicate the findings from Experiment I and additionally validate the UCS effects in an assay specifically suited to detect anxiety-like behavior. As in Experiment I, animals were first assessed in the SPT or SCT, and the same individuals were submitted to the NTT on the following day. This approach allowed a comparison of social and anxiety-like responses within the same animals, enabling us to evaluate whether UCS selectively affected exploration in a novel environment while leaving social behavior intact.

**Figure 1.**
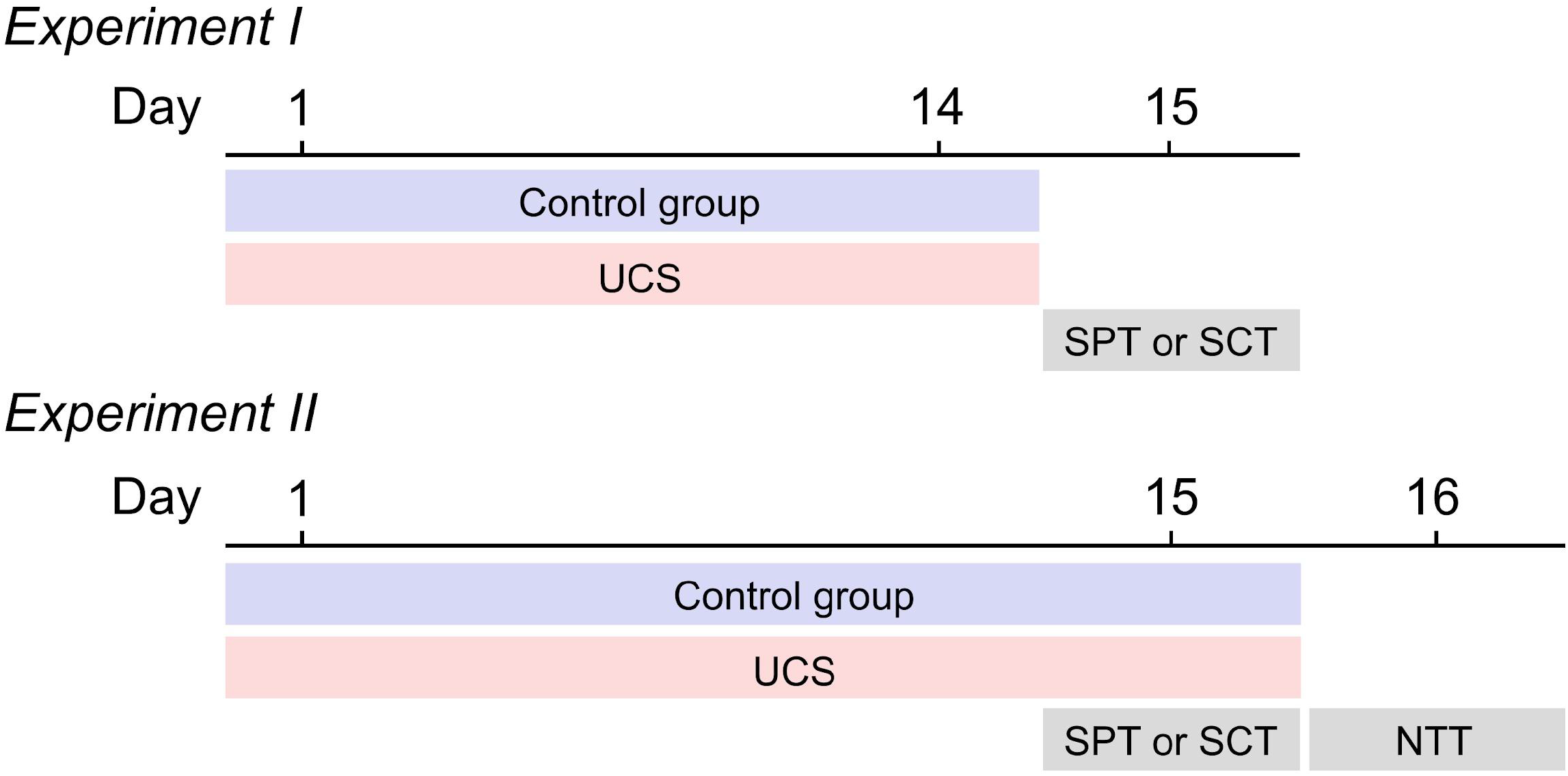
Overview of the experimental design evaluating the effects of unpredictable chronic stress (UCS) on the behavior of adult zebrafish. SPT: social preference test; SCT: shoal cohesion test; NTT: novel tank test.

### Unpredictable chronic stress protocol (UCS)

After the two-week acclimation, fish were assigned to experimental groups using block randomization stratified by sex and original housing tank. Within each block, individuals were allocated to new experimental tanks according to an independently generated random sequence from an online randomizer (random.org). Animals were distributed to either (i) the control group, kept undisturbed except for routine husbandry (feeding and minimal handling during necessary tank cleaning), or (ii) the unpredictable chronic stress (UCS) group, which underwent a 14-day UCS protocol as described in detail elsewhere [13].

Across 14 days, fish in the UCS group experienced two stressors per day: one in the morning and a different one in the afternoon. The type of stressor and start time varied daily to reduce predictability (windows approximately 08:00 a.m. to 12:00 p.m. and 01:00 p.m. to 06:00 p.m.). The stressor set comprised: (1) heating the tank to 33 °C for 30 min; (2) cooling to 23 °C for 30 min; (3) crowding 12 fish in a 600 mL beaker containing 250 mL of water for 50 min; (4) lowering the water level until dorsal exposure for 2 min; (5) three sequential tank transfers for 10 min each in tanks of different dimensions; and (6) chase with a net for 8 min.

After completing the 14-day protocol, fish were tested in the SPT and SCT the next morning (experimental day 15). In Experiment II, the NTT was conducted the following morning (experimental day 16), and a randomly selected additional stressor (chasing with a net in this study) was applied in the afternoon of day 15.

### Behavioral analysis

Within each experiment, all SPT trials were completed first, followed by all SCT sessions. Behavioral testing took place during the light phase, between 07:00 am and 12:00 pm. The testing sequence was block-randomized by housing tank and test arena, and across blocks the run order was generated using random.org. Behavioral data for SPT and NTT were analyzed using ANY-maze (version 7.20). SCT videos were scored in ImageJ (version 1.54). All data extraction and scoring were performed by researchers blinded to the experimental groups.

### Social preference test (SPT)

The SPT was used to assess social approach, as described previously [14,15]. Each zebrafish was placed individually in a central tank (30 × 10 × 15 cm) flanked by two side tanks (15 × 10 × 13 cm). One tank contained 10 conspecific zebrafish (social stimulus), and the other only water (neutral stimulus). All tanks were filled with dechlorinated water to a depth of 10 cm. The position of the social stimulus tank (left or right) was alternated between trials to prevent bias, and the water in the central tank was replaced after each trial to eliminate olfactory cues. Social behavior was recorded for 7 minutes: 2 minutes for habituation, followed by 5 minutes for behavioral evaluation. The central tank was divided into three vertical zones: the interaction zone (next to the social stimulus), the intermediary zone (center), and the neutral zone (next to the neutral stimulus). In this task, greater time spent in the interaction zone reflects increased social preference, whereas reduced time in this zone indicates diminished social approach. We quantified time spent in the interaction zone, total distance traveled, number of zone crossings, maximum speed, and immobility time.

### Shoal cohesion test (SCT)

The SCT was adapted from a previously described protocol [16]. Groups of four zebrafish were transferred to a 2.7-L tank (24 × 8 × 20 cm), which provides a controlled environment that allows detailed observation of spacing and the overall structure of a small shoal. Using four fish facilitates the measurement of cohesion and individual positioning without visual overlap between animals, making it well suited for quantitative analyses. In this assay, higher cohesion is reflected by fish maintaining closer distances and a more compact group structure, whereas increased distances indicate reduced shoal cohesion. Each group was recorded continuously for one hour. After recording, screenshots were extracted every 15 seconds within three predefined 5-minute intervals (0–5 min, 30–35 min, and 55–60 min). The images were analyzed in ImageJ using the Interfish Distance plugin, as described in a step-by-step protocol [17]. For each frame, a point was placed at the center of each fish, allowing calculation of the interfish distance, nearest neighbor distance, and farthest neighbor distance. Shoal area and perimeter were obtained by outlining the convex hull of the group, such that the innermost fish was excluded from the outline when group geometry would yield a concave quadrilateral. All measurements were averaged per shoal, which served as the experimental unit.

### Novel tank test (NTT)

The NTT, performed as described elsewhere [18], was used to assess exploratory patterns and anxiety-like behavior in a novel environment. The assay was conducted in a 2.7-L tank (24 × 8 × 20 cm) filled with dechlorinated water to a depth of 18 cm and virtually divided into three equal horizontal zones. Each fish was individually introduced into the tank and recorded for 6 minutes. Video recordings were analyzed to obtain the average height in the tank, time spent in the upper zone, number of entries into the upper zone, total distance traveled, number of zone crossings, and time spent immobile. Average height in the tank corresponds to the mean vertical position (y-coordinate) of the fish in the water column, measured from the bottom of the apparatus.

### Statistical analysis

Sample size estimation was performed for the SPT based on the effect of UCS on social behavior reported by a previous study [19]. A priori calculations were performed in G*Power 3.1 using the following parameters: effect size = 1.19, α = 0.05, and power = 0.95. This resulted in a required sample of 20 animals per group for the primary outcome (time spent in the interaction zone). Accordingly, each batch of the SPT included 20 animals per group. For the shoal cohesion test, sample size was adapted to the constraints of available arenas and the one-hour duration of each trial; with eight test arenas, four shoals per group were included in each batch. Experiment I comprised two batches (data were pooled), while Experiment II comprised only one batch. Detailed sample sizes for each assay and experiment are described in Table 1.

**Table 1.** Effect sizes (Cohen’s d) and 95% confidence intervals for the effects of unpredictable chronic stress (UCS) on all behavioral outcomes across experiments.

| Test / Experiment | Outcome | Cohen's d [CI lower, CI upper] | N UCS | N CTRL |
| --- | --- | --- | --- | --- |
| Social Preference – Exp. I | Interaction time | 0.161 [-0.279, 0.599] | 40 | 40 |
|  | Distance | -0.419 [-0.860, 0.026] | 40 | 40 |
|  | Time immobile | -0.426 [-0.868, 0.019] | 40 | 40 |
|  | Line crossings | -0.191 [-0.629, 0.249] | 40 | 40 |
|  | Max speed | -0.094 [-0.532, 0.345] | 40 | 40 |
| Shoal Cohesion – Exp. I | Interfish distance | 0.260 [-0.730, 1.240] | 8 | 8 |
|  | Nearest neighbor | 0.131 [-0.853, 1.109] | 8 | 8 |
|  | Farthest neighbor | 0.305 [-0.686, 1.286] | 8 | 8 |
|  | Area | 0.007 [-0.973, 0.987] | 8 | 8 |
|  | Perimeter | 0.211 [-0.776, 1.191] | 8 | 8 |
| Social Preference – Exp. II | Interaction time | -0.177 [-0.797, 0.445] | 20 | 20 |
|  | Distance | 0.025 [-0.595, 0.645] | 20 | 20 |
|  | Time immobile | -0.204 [-0.824, 0.419] | 20 | 20 |
|  | Line crossings | 0.200 [-0.423, 0.820] | 20 | 20 |
|  | Max speed | -0.005 [-0.625, 0.614] | 20 | 20 |
| Shoal Cohesion – Exp. II | Interfish distance | 1.021 [-0.511, 2.482] | 4 | 4 |
|  | Nearest neighbor | 0.955 [-0.562, 2.404] | 4 | 4 |
|  | Farthest neighbor | 1.054 [-0.486, 2.521] | 4 | 4 |
|  | Area | 0.686 [-0.776, 2.097] | 4 | 4 |
|  | Perimeter | 1.036 [-0.499, 2.500] | 4 | 4 |
| Novel Tank – Exp. II | Average height | -0.980 [-1.466, -0.487] | 36 | 36 |
|  | Upper time | -0.823 [-1.302, -0.339] | 36 | 36 |
|  | Upper entries | -0.478 [-0.945, -0.007] | 36 | 36 |
|  | Distance | -0.431 [-0.897, 0.038] | 36 | 36 |
|  | Line crossings | -0.540 [-1.009, -0.068] | 36 | 36 |
|  | Time immobile | 0.265 [-0.200, 0.728] | 36 | 36 |

Effect sizes are reported for each outcome in the Social Preference Test and Shoal Cohesion Test in Experiment 1, and for the corresponding outcomes in Experiment 2, including the Novel Tank Test. Values are presented as the estimated effect size followed by the lower and upper limits of the confidence interval. N UCS and N CTRL denote the number of animals or shoals included in each comparison. CTRL: control group.

Statistical analyses were conducted in GraphPad Prism (version 10.1.0). To adopt a conservative approach and because assumptions of normality were not consistently met across datasets, comparisons between groups were performed using the nonparametric Mann–Whitney test. Results are presented as median ± interquartile range, and p-values below 0.05 were considered statistically significant. Effect sizes (Cohen’s d) and 95% confidence intervals were calculated in R (version 4.3.1) using RStudio (version 2025.09.2) and the “effectsize” package [20]. Raw data and data supporting the analyses are available via the Open Science Framework [12].

The experiments conducted here were neither designed nor powered to detect the effects of sex or time (e.g., temporal bins in the SCT). Consequently, behavioral data presented in the main manuscript are pooled across sex and, for the SCT, across time intervals. Supplementary material provides descriptive, sex-disaggregated plots for the SPT and NTT, as well as graphical representations of SCT outcomes across time points; no inferential statistics were performed on these datasets due to insufficient statistical power and because such analyses fell outside the scope of the study design.

## RESULTS

### Experiment I

Regarding the SPT, UCS did not change social approach, as indicated by the absence of group differences in time spent in the interaction zone (Figure 2A; U = 726, p = 0.4800), total distance traveled (Figure 2B; U = 627.5, p = 0.0976), number of line crossings (Figure 2C; U = 756, p = 0.6754), maximum speed (Figure 2D; U = 607.5, p = 0.0641) or immobility time (Figure 2E; U = 670, p = 0.1991). All effect sizes had confidence intervals that crossed zero (Table 1), consistent with no differences between groups.

**Figure 2.**
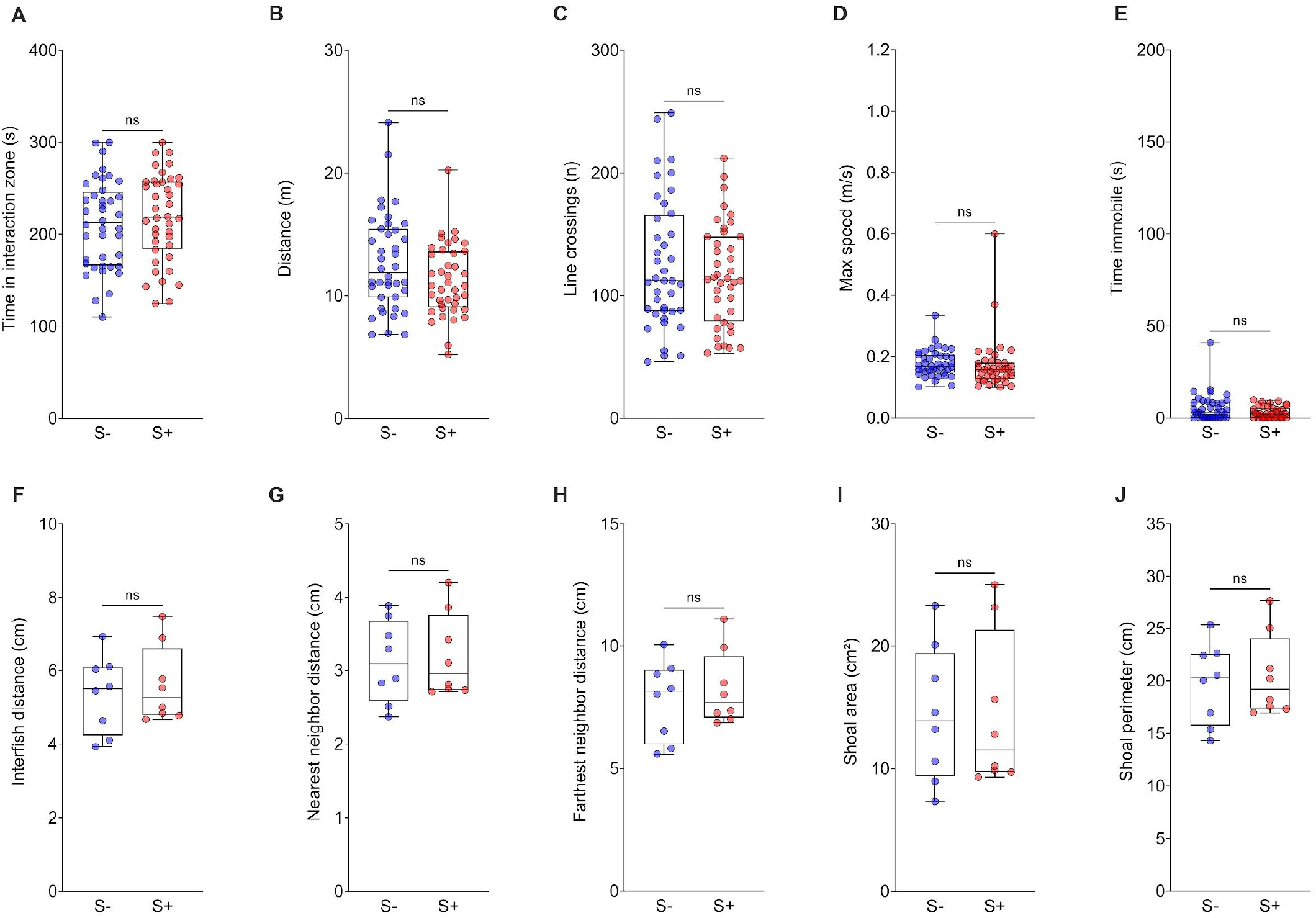
Effects of unpredictable chronic stress (UCS) on the behavior of adult zebrafish in the social preference test (A–E) and shoal cohesion test (F–J) in Experiment I. (A) Time spent in the interaction zone, (B) total distance traveled, (C) number of line crossings, (D) maximum swimming speed, and (E) time spent immobile (n = 40). (F) Mean interfish distance, (G) mean nearest neighbor distance, (H) mean farthest neighbor distance, (I) shoal area, and (J) shoal perimeter (n = 8 shoals per group). Data are shown as median with interquartile range. Group comparisons were performed using the Mann–Whitney test. S-: control group; S+: UCS-exposed group.

Regarding group-level social behavior, none of the cohesion metrics differed between UCS-exposed and control shoals: interfish distance (Figure 2F; U = 28, p = 0.7209), nearest neighbor distance (Figure 2G; U = 32, p > 0.999), farthest neighbor distance (Figure 2H; U = 29, p = 0.7984), shoal area (Figure 2I; U = 31, p = 0.9591) or shoal perimeter (Figure 2J; U = 28, p = 0.7209). Effect sizes similarly showed confidence intervals spanning zero (Table 1), further supporting the absence of UCS effects on shoal cohesion.

Sex-disaggregated plots in Supplementary Figure 1 (A–E) revealed similar patterns for females and males, with no clear sex-specific effects upon visual inspection. Supplementary Figure 2 (A–E) shows the progression of shoal cohesion across the three time intervals. All measures increased over time in both groups, consistent with a gradual reduction in cohesion as fish habituated to the tank and increased exploration. No apparent differences were observed between UCS and control shoals.

### Experiment II

Replicating the findings from Experiment I, UCS did not affect time spent in the interaction zone (Figure 3A; U = 191, p = 0.8196), total distance traveled (Figure 3B; U = 183, p = 0.6588), line crossings (Figure 3C; U = 193, p = 0.8562), maximum speed (Figure 3D; U = 164.5, p = 0.3441), or immobility time (Figure 3E; U = 171, p = 0.3521) in the SPT. All effect size confidence intervals included zero (Table 1), indicating no significant difference between UCS and control fish.

**Figure 3.**
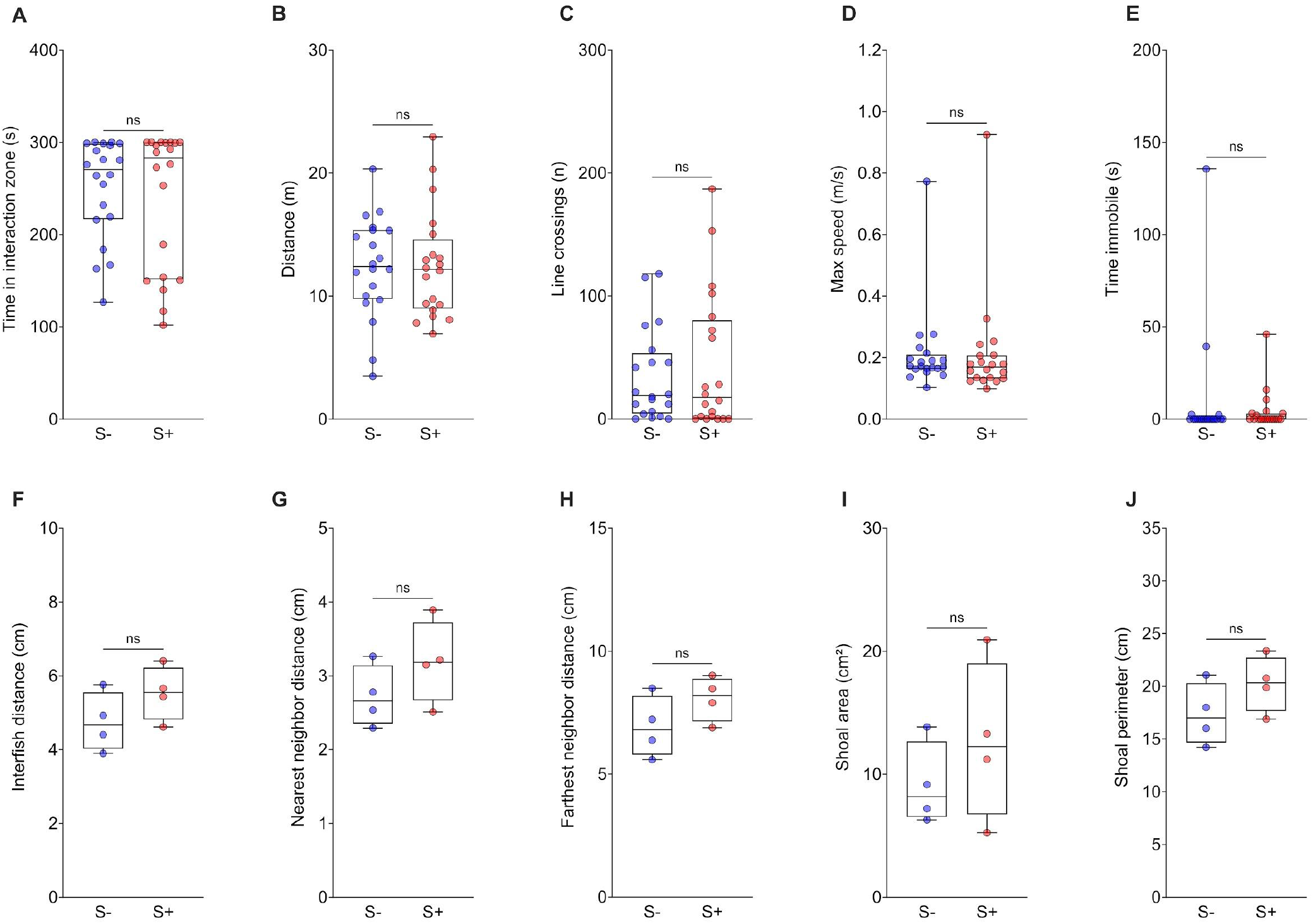
Effects of unpredictable chronic stress (UCS) on the behavior of adult zebrafish in the social preference test (A–E) and shoal cohesion test (F–J) in Experiment II. (A) Time spent in the interaction zone, (B) total distance traveled, (C) number of line crossings, (D) maximum swimming speed, and (E) time spent immobile (n = 20). (F) Mean interfish distance, (G) mean nearest-neighbor distance, (H) mean farthest-neighbor distance, (I) shoal area, and (J) shoal perimeter (n = 4 shoals per group). Data are shown as median with interquartile range. Group comparisons were performed using the Mann–Whitney test. S-: control group; S+: UCS-exposed group.

Similarly, UCS again did not alter shoal cohesion: interfish distance (Figure 3F; U = 4, p = 0.3429), nearest neighbor distance (Figure 3G; U = 5, p = 0.4857), farthest neighbor distance (Figure 3H; U = 4, p = 0.3429), shoal area (Figure 3I; U = 6, p = 0.6857), or shoal perimeter (Figure 3J; U = 4, p = 0.3429). Confidence intervals for all effect sizes crossed zero (Table 1).

Across the SCT time intervals, all cohesion measures gradually increased for both groups (Supplementary Figure 2 F–J), reflecting reduced shoal cohesion with habituation. This temporal progression was comparable between UCS-exposed and control fish.

These findings corroborate Experiment I and support the interpretation that social behavior in zebrafish remains robust, showing no evidence of alteration following UCS exposure.

In contrast to the social assays, UCS produced clear anxiety-like behavior in the NTT. UCS-exposed fish swam at lower average heights (Figure 4A; U = 291.5, p < 0.0001), spent less time in the upper zone (Figure 4B; U = 329, p = 0.0002), and made fewer entries into the upper zone (Figure 4C; U = 425, p = 0.0114). Line crossings were also reduced (Figure 4E; U = 445, p = 0.0217). Total distance traveled (Figure 4D; U = 515, p = 0.1363) and immobility time (Figure 4F; U = 593, p = 0.3969) did not differ. Effect sizes for vertical exploration measures were moderate to large, and their confidence intervals did not include zero (Table 1), confirming a consistent anxiogenic effect of UCS and validating the study. Sex-disaggregated plots (Supplementary Figure 3) showed the same pattern in males and females.

**Figure 4.**
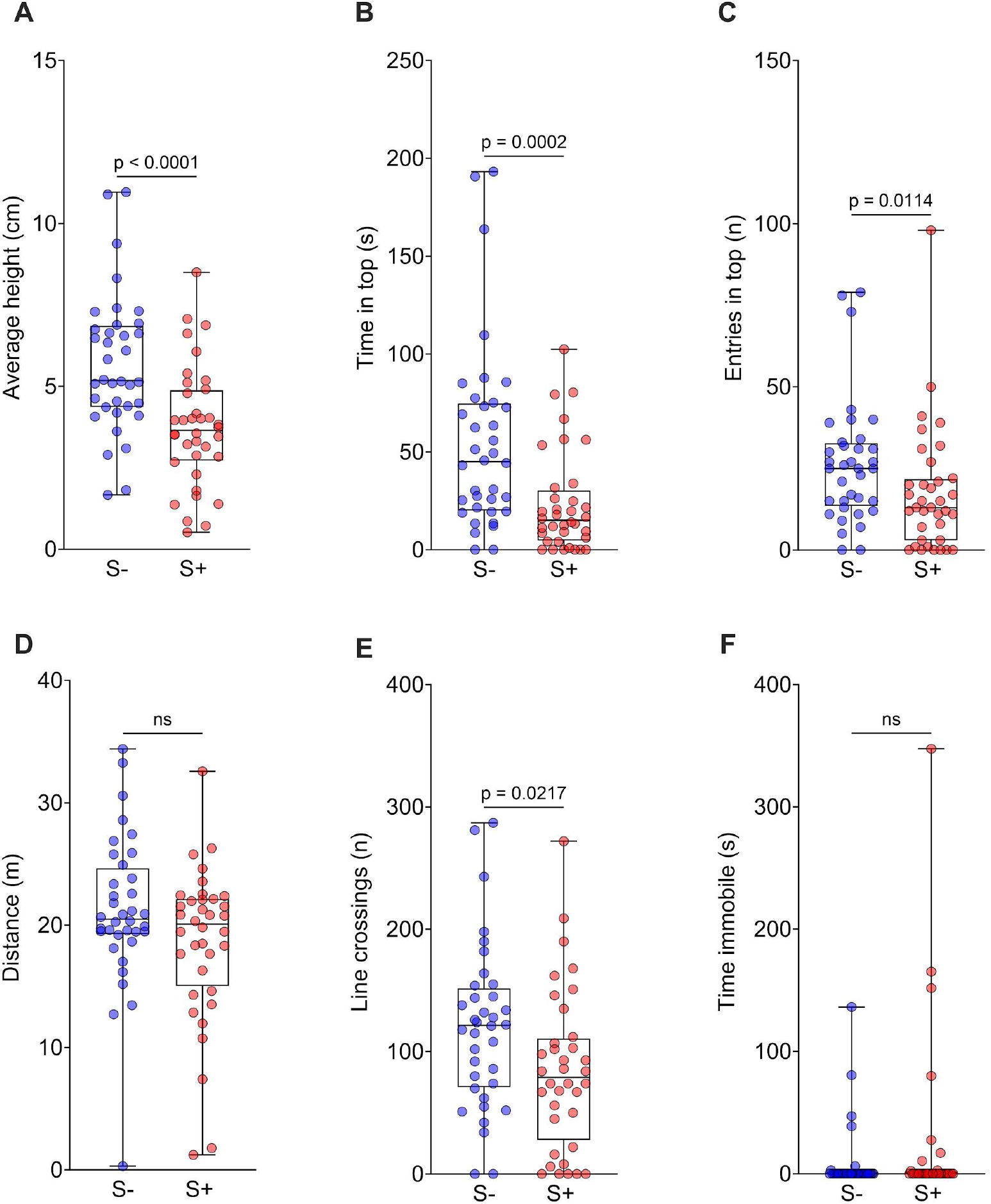
Effects of unpredictable chronic stress (UCS) on the behavior of adult zebrafish in the novel tank test in Experiment II. (A) Average height, (B) time spent in the upper zone, (C) entries in the upper zone, (D) total distance traveled, (E) line crossings, and (F) time spent immobile. (n = 36). Data are presented as median with interquartile range. Group comparisons were performed using the Mann–Whitney test. S-: control group, S+: UCS-exposed group.

## DISCUSSION

A decade-wide systematic review and meta-analysis of UCS studies in zebrafish reported that, although the paradigm reliably increases anxiety-like behavior and elevates cortisol levels, its effects on social behavior are highly inconsistent, and the pooled effect size for social outcomes was non-significant [11]. The present findings align with the meta-analytic summary: across two independent experiments and two complementary assays, the UCS protocol did not alter social preference or shoal cohesion in adult zebrafish. These data add to the limited number of well-controlled studies examining stress-induced social outcomes and support the conclusion that UCS does not reliably impair sociability in this species.

The previous studies reported highly divergent effects of UCS on zebrafish social behavior. Some studies describe increased shoal cohesion following repeated stress [21,22], whereas others document reduced social preference or disrupted shoaling [10,23,24]. A third group of studies finds no observable change in social behavior [25–27]. Such discrepancies likely reflect methodological variation, including differences in stressor intensity and duration, developmental stage, genetic background, housing density, and the specific behavioral paradigms used to quantify social behavior. In addition, these inconsistencies may reflect the cumulative impact of study-specific biases and statistical limitations, including limited sample sizes, low statistical power, and an increased likelihood of false-positive findings. Our findings contribute to clarifying this landscape by demonstrating, under controlled conditions and with appropriate sample sizes, that UCS does not impair either individual social approach or group-level cohesion.

One interpretation is that social behavior in adult zebrafish is relatively resilient to chronic stress [11]. Social interaction is a core ethological drive in this species [28–30]. Zebrafish display robust social preference early in development and form cohesive shoals as adults, a behavior that improves predator evasion and foraging efficiency [31–33]. At the neural level, social behavior is supported by a conserved subcortical network, and recent whole-brain activity mapping indicates the existence of dedicated ensembles encoding individual recognition and long-term social memory [34,35]. Because social behavior is strongly conserved and tightly regulated in zebrafish, it may be less sensitive to perturbations such as those imposed by UCS. In this context, the stress protocol used here, while sufficient to induce clear anxiogenic effects, may not have reached the threshold needed to disrupt strongly conserved social drives.

Our results also showed that shoal cohesion decreased over time during the SCT in both control and UCS groups, which is consistent with habituation to the test environment. As zebrafish become familiar with the arena, they typically increase exploratory behavior and reduce group cohesion. Crucially, this temporal pattern was indistinguishable between experimental groups, reinforcing the notion that UCS did not alter social organization at any point during the assay. This finding supports the idea that shoaling patterns are driven more by acclimation to novelty than by the prior stress history of the fish under the conditions tested.

While UCS did not disrupt social behavior, it produced a clear anxiogenic phenotype in the NTT. Stressed fish showed reduced vertical exploration, decreased time and entries into the upper zone, and lower average swimming height, all hallmark indices of heightened anxiety-like behavior in zebrafish [36–38]. Importantly, these effects were observed in the same animals that exhibited intact social preference and shoal cohesion. This dissociation suggests that UCS may selectively affect behavioral domains differentially sensitive to stress, influencing responses related to novelty and risk assessment while sparing affiliative behaviors.

Several methodological considerations warrant discussion. First, although we formed shoals randomly to avoid selection bias, this also meant that sex composition of shoals varied, which may introduce subtle variability. However, the lack of sex effects in the supplementary analyses suggests that this factor likely did not influence our conclusions. Second, although the UCS protocol used here is widely adopted, different durations or intensities of stress might yield other outcomes. Third, as social hierarchy can influence zebrafish social dynamics and stress reactivity, future studies incorporating dominance assessments may help clarify whether hierarchical structure modulates stress effects on sociability.

Collectively, the present findings contribute to a clearer understanding of how unpredictable chronic stress shapes zebrafish behavior. We show that while UCS produces robust anxiety-like behavior, it does not impair individual social preference or shoal cohesion in adult zebrafish. These findings support the idea that social behavior in zebrafish may be relatively resistant to chronic stress under many experimental conditions. Future work should systematically examine how factors such as stress intensity, developmental stage, sex composition, social rank, and assay-specific features interact to determine whether and when UCS affects social outcomes.

## AUTHOR CONTRIBUTIONS

Conceptualization: N.A.N., M.G.-L., A.P.H. and A.P. Data curation: M.G.-L., A.P.-A. and D.V.M. Formal analysis: M.G.-L., A.P.-A. Funding acquisition: A.P. Investigation: N.A.N., M.G.-L., A.P.-A., D.V.M., L.M.B. and T.S.-B. Methodology: N.A.N., M.G.-L., A.P.H. and A.P. Project administration: A.P. Supervision: A.P. Writing - original draft: N.A.N., M.G.-L., A.P.-A., D.V.M. and A.P. Writing - review & editing: L.M.B., T.S.-B. and A.P.H..

## DATA AVAILABILITY

All data supporting this study are publicly accessible via the Open Science Framework (osf.io/u8pkm) [12].

## ACKNOWLEDGMENTS

We thank Conselho Nacional de Desenvolvimento Científico e Tecnológico (CNPq, grant number 305968/2023-8), Coordenação de Aperfeiçoamento de Pessoal de Nível Superior - Brazil (CAPES), and Pró-Reitoria de Pesquisa (PROPESQ) at Universidade Federal do Rio Grande do Sul (UFRGS) for funding and support.

## DECLARATION OF GENERATIVE AI AND AI-ASSISTED TECHNOLOGIES IN THE MANUSCRIPT PREPARATION PROCESS

ChatGPT-5.1 was used to help refine wording and formatting of the manuscript. All outputs were reviewed and amended by the authors, who take full responsibility for the final text and its interpretation.

