## Supplementary Files for "Unpredictable chronic stress does not alter social behavior in zebrafish"

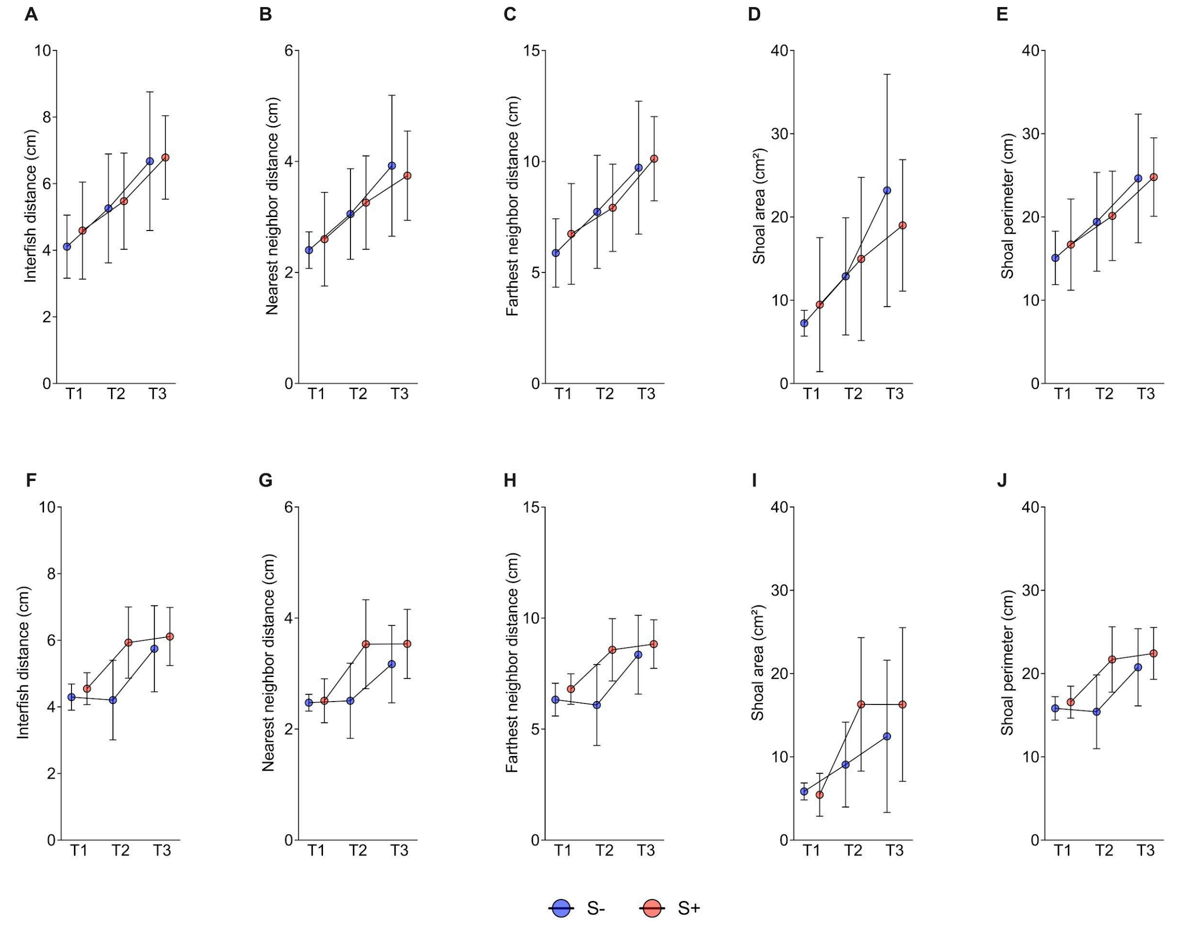


**Supplementary Figure 1.** Effects of unpredictable chronic stress (UCS) on the behavior of female and male adult zebrafish in the social preference test in Experiment I. (A) Time spent in the interaction zone, (B) total distance traveled, (C) number of line crossings, (D) maximum swimming speed and (E) time spent immobile. Data are presented as median with interquartile range. Sample sizes: S-: F = 13, M = 27; S+: F = 20, M = 20.

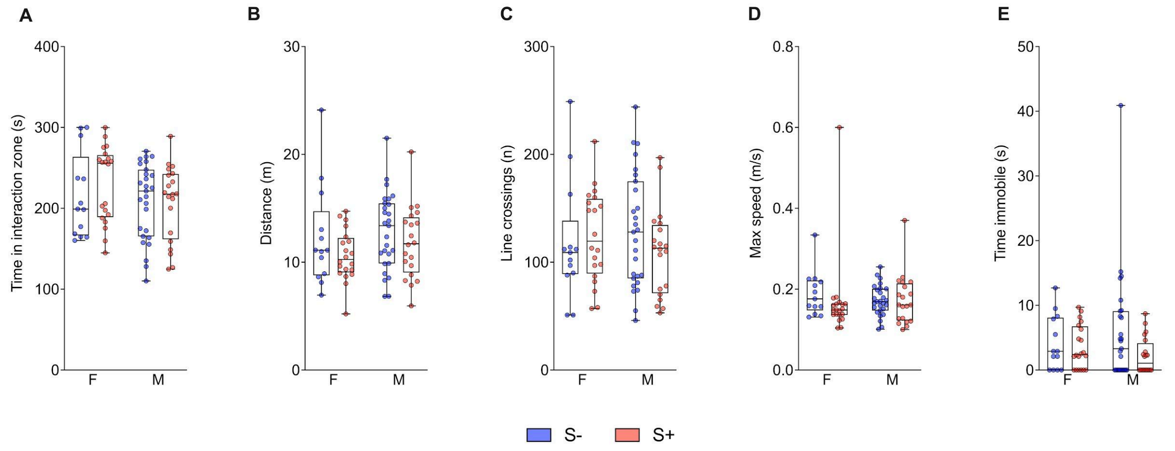


**Supplementary Figure 2.** Effects of unpredictable chronic stress (UCS) on shoal cohesion test in Experiment I (A-E), and in Experiment II (F-J). Shoal cohesion was quantified at three time intervals: T1 (0–5 min), T2 (30–35 min), and T3 (55–60 min). (A) Mean interfish distance, (B) mean nearest neighbor distance, (C) mean farthest neighbor distance, (D) shoal area and (E) shoal perimeter. (n = 8 shoals per group). (F) Mean interfish distance, (G) mean nearest neighbor distance, (H) mean farthest neighbor distance, (I) shoal area and (J) shoal perimeter. Data are presented as mean ± S.D.

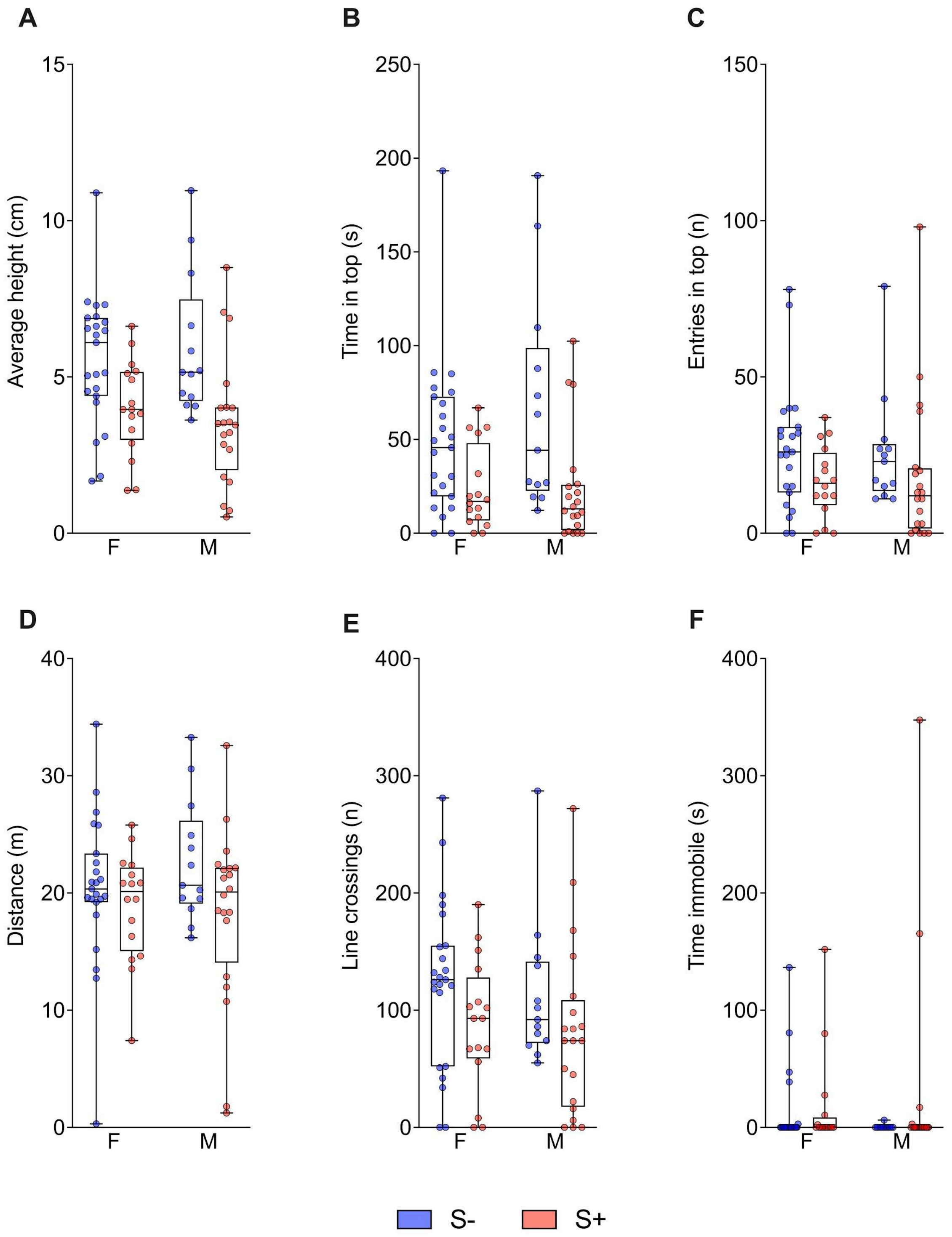


**Supplementary Figure 3.** Effects of unpredictable chronic stress (UCS) on the behavior of female and male adult zebrafish in the novel tank test. (A) Average height, (B) time spent in the upper zone, (C) entries in the upper zone, (D) total distance traveled, (E) line crossings, and (F) time spent immobile. Data are presented as median with interquartile range. F: female, M: male, S-: control group, S+: UCS-exposed group. Sample sizes: S-: F= 23, M= 13; S+: F = 16, M = 20.
